# Comparative *in silico* analysis of *ftsZ* gene from different bacteria reveals the preference for core set of codons in coding sequence structuring and secondary structural elements determination

**DOI:** 10.1101/676932

**Authors:** Ayon Pal, Barnan Kr Saha, Jayanti Saha

## Abstract

The deluge of sequence information in the recent times provide us with an excellent opportunity to compare organisms on a large genomic scale. In this study we have tried to decipher the variation in the gene organization and structuring of a vital bacterial gene called *ftsZ* which codes for an integral component of the bacterial cell division, the FtsZ protein. FtsZ is homologous to tubulin protein and has been found to be ubiquitous in eubacteria. FtsZ is showing increasing promise as a target for antibacterial drug discovery. Our study of *ftsZ* protein from 143 different bacterial species spanning a wider range of morphological and physiological type demonstrates that the *ftsZ* gene of about ninety three percent of the organisms involved in our analyses show relatively biased codon usage profile and significant GC deviation from their genomic GC content. We have also detected a tendency among the different organisms to utilize a core set of codons in structuring the *ftsZ* coding sequence. Our meticulous analysis of the *ftsZ* gene linked with the corresponding FtsZ protein show that there is a bias towards the use of specific synonymous codons particularly in the helix and strand regions of the multi-domain FtsZ protein. Overall our findings suggest that in an indispensable and vital protein such as FtsZ, there is an inherent tendency to maintain form and structure for optimized performance in spite of the extrinsic variability in coding features.

## Introduction

Codon usage bias (CUB) or the preference of an organism for a certain subset of codons coding for the different amino acids of polypeptides has intrigued molecular biologists and evolutionists for decades [1]. This is a universal phenomenon observed in prokaryotes, eukaryotes [2] as well as viruses [3] and is predominantly dependent on selection, mutation, and genetic drift [4]. CUB has been found to be an important factor contributing to gene and genome evolution [5, 6] and has also been found to be an important determinant of gene expression levels at the transcription level [2]. Codon usage pattern has not only been found to vary between organisms but also between coding sequences or genes within an organism [4]. In this study we have tried to decipher the variation in the gene organization and structuring of a vital bacterial gene called *ftsZ* which codes for an integral component of the bacterial cell division, the FtsZ protein. The process of bacterial cytokinesis is initiated by the assembly of the tubulin-like GTPase called FtsZ which is essential for bacterial cell division [7]. FtsZ is homologous to tubulin protein which acts as the building block of the microtubule cytoskeleton in eukaryotes FtsZ [8]. During cell division, FtsZ interacts with other membrane associated proteins like FtsW, FtsK, FtsQ and FtsI and helps in anchoring FtsZ to the bacterial cytoplasmic membrane [9]. FtsZ is reported to be a highly conserved protein [8] with a relative molecular mass of 40,000 and is ubiquitous in eubacteria. It is also found in the members of Euryarchaea, chloroplasts of plants and some mitochondria [10]. Higher plants have also been found to contain two distinct families of FtsZ homologues that seem to have diverged early in the evolution of plants [11]. Mutant bacteria which lacks FtsZ protein cannot divide but elongate into filamentous form. During cytokinesis, the FtsZ protein assemble into a contractile ring that provides a stage for assembly of the cell division apparatus and constricts at the leading edge of the invaginating septum [7]. FtsZ is a vital cell-division protein in prokaryotes and is showing increasing promise as a target for antibacterial drug discovery [12]. Looking at the ubiquity and conserved nature of FtsZ, it has been projected as a potent target and has been studied extensively [13] for the discovery of next-generation antibacterial agents that can be used to counter drug-resistances to the commonly used drugs for methicillin resistant *Staphylococcus aureus* (MRSA), tuberculosis, and other microorganism mediated infections [14]. The *ftsZ* gene is regarded as an essential cell division gene in many bacteria including *E. coli* [15] and it has been found that the C-terminal domain for FtsZ is highly variable in both size and alignment among the bacterial species [16].

The major objectives of our study was to decipher the codon usage pattern of the *ftsZ* gene to find out if there exist any codon usage bias in the structuring of the *ftsZ* coding sequence among different types of bacteria. We have tried to find out if the codon usage pattern is a random phenomenon or has it been influenced by certain features such as the lifestyle of the organism [17–21]. This includes their free-living behaviour or pathogenic association with specific host organisms and ecological associations. We have also tried to unravel whether the codon structuring of the *ftsZ* gene is to certain extent influenced by the Gram nature of the organism. The Gram nature of a bacterium, although primarily attributed to the cell wall construction of the bacterium, has been found to manifest a host of comparative features in the organisms ranging from simple morphology to advanced physiological, biochemical, ecological and molecular characteristics such as GC content. We have also attempted to estimate the compositional divergence of the *ftsZ* coding sequences. The FtsZ protein is a very vital component of bacterial cell division that demonstrates promiscuous variability both in terms of gene sequence and amino acid composition. This compositional variability in a conserved protein such as FtsZ has been our impetus to decipher and track whether there exists the preference for a ‘core’ set of codons in coding the gene sequence across a diverse group of bacteria. In our study we have tried to explore the codon usage tendency based on the positioning of the different amino acids in the different types of structural elements of the FtsZ protein. It has been reported that codon usage can play an important role in the translation process as well as the folding behaviour of nascent polypeptides [22, 23]. We have adopted a unique approach to further explore the codon usage bias profile of the *ftsZ* sequence by linking the codon utilization profile with the secondary structural components of the protein. Thus, we have strived to correlate the coding pattern of the *ftsZ* gene with the structural attributes of the FtsZ protein. We have meticulously analysed the 61 sense codons coding for the twenty standard amino acids to find out the preference of disposition of specific codons in specific secondary structural elements of the FtsZ protein.

## Materials and methods

The *ftsZ* gene sequence of 143 bacteria were selected, and their whole genome sequences were retrieved from the NCBI GenBank [24] sequence database. The *ftsZ* coding sequences (CDS) and their corresponding amino acid sequences were screened out from the whole genome sequences of the bacteria using Perl scripts generated in our lab. Analysis of different codon usage bias parameters like effective number of codons (Nc) [25], GC content, guanine and cytosine content at the third position of the codon (GC3) [25] and hydrophobicity were also estimated.

The Nc determines the degree of bias for the use of codons [26] with value ranging from 20 to 61, where lower value indicates higher codon usage bias and vice versa. The GC content plays a critical role in genome evolution [27], and it has been found to range from 13% to 75% in cellular organisms [28, 29]. The GC content does not remain constant throughout the genome of an organism but varies based on different regions and coding sequences of the genome. The measurement of different GC based attributes like GC content and GC3 content thus play a significant role in analysing the genomic as well as genic organization. The GC3 and GC content of each individual *ftsZ* sequence was calculated using our in house developed tool using Perl. The Nc–plot [25], which is a parabolic curve used to measure and explore codon usage bias, and detect the effect of base content on CUB [30] was also constructed.

Statistical analysis such as non-parametric One way ANOVA on Ranks [31] was used to find out whether there is a preferred set of codon for each of the amino acid that is used in the structuring of the *ftsZ* coding sequences. Two factor ANOVA on codon usage of *ftsZ* CDS was also performed to study the frequency of the individual 61 sense codons and their interrelation with lifestyle and Gram nature of the organisms. A two factor ANOVA was also designed to study the interrelationship of the twenty different amino acids with lifestyle and Gram nature of the bacteria.

The degree of identity in FtsZ protein sequences among the 142 organisms considered for this study was analysed using Clustal Omega. This application employs HMM profile-profile techniques along with seeded guide trees to produce multiple alignments [32]. For clustering of similar proteins based on their sequence similarities, the program CD-HIT [33] was used. All the 143 *ftsZ* CDS were subjected to clustering using CD-HIT with a similarity threshold of 50%. Representative amino acid sequences of *ftsZ* of the four main clusters as identified by CD-HIT were subjected to secondary structure (helix, strands and other elements) prediction using SSpro module of SCRATCH Protein Predictor [34]. Accurately predicting protein secondary structure is important for the study of protein evolution, structure and function. The SSpro program was accessed through the SCRATCH suite of protein structure predictors hosted at http://scratch.proteomics.ics.uci.edu.

The *ftsZ* gene sequences were further aligned with their corresponding amino acid sequences and secondary structure mark-up sequence generated using SSpro. With the help of this triple alignment, we have identified each of the synonymous codons that are used for coding the amino acids, and we have linked those codons with the amino acids of the predicted secondary structural elements. The relative synonymous codon usage (RSCU) value which is measured by the ratio between the actual observed values of the codon and the theoretical expectations was also calculated. RSCU reflects the relative usage preference for the specific codons encoding the same amino acid [35]. If RSCU value equals to 1, codon usage is supposed to be unbiased but if RSCU>1, specific codon frequency is higher than other synonymous codons and codon usage is considered to be biased [26]. The RSCU values of the *ftsZ* CDS were calculated after splitting the sequences based on their propensity in constituting the different secondary structural classes as predicted by SSpro.

## Results and discussion

A comprehensive codon usage analysis of the *ftsZ* gene and its corresponding protein (FtsZ) was carried out in 143 spp. of bacteria of which 74 are non-pathogenic and 69 are pathogenic in nature. On the basis of the nature of cell wall, 43 are Gram positive, 99 organisms are Gram negative and one organism called *Gardnerella vaginalis* 409-05 is Gram variable in nature. A list of the organisms considered in this study along with their Gram nature and lifestyle is presented in Table 1.

**Table 1:** Details of the different bacterial species along with their lifestyle, Gram nature and codon usage attributes considered in the study.

| Organism | Lifestyle | Gram Nature | Genomic Nc | *ftsZ* Nc | Genomic GC3 | *ftsZ* GC3 | Genomic GC | *ftsZ* GC |
| --- | --- | --- | --- | --- | --- | --- | --- | --- |
| *Acetobacter malorum* | NP | Negative | 45.41 | 40.62 | 0.69 | 0.6933 | 56.5 | 62.62 |
| *Acinetobacter baumannii* | P | Negative | 46.408 | 40.11 | 0.2974 | 0.1885 | 39 | 43.11 |
| *Acinetobacter johnsonii* XBB1 | NP | Negative | 47.345 | 39.98 | 0.39 | 0.3384 | 38.5 | 45.95 |
| *Actinobacillus pleuropneumoniae* serovar 5b str. L20 | P | Negative | 43.98 | 42.48 | 0.48 | 0.4796 | 41.3 | 47.11 |
| *Actinomyces odontolyticus* ATCC 17982 | P | Positive | 37.24 | 33.9 | 0.808 | 0.7556 | 65.4 | 66.67 |
| *Aerococcus viridans* | P | Positive | 47.09 | 34.65 | 0.339 | 0.1463 | 39.4 | 43.81 |
| *Aeromonas enteropelogenes* | P | Negative | 39.009 | 31.64 | 0.821 | 0.7818 | 60 | 62.24 |
| *Afipia broomeae* ATCC 49717 | NP | Negative | 42.26 | 34.41 | 0.8 | 0.8211 | 61.3 | 68.2 |
| *Aggregatibacter actinomycetemcomitans* | P | Negative | 45.48 | 43.73 | 0.56 | 0.4048 | 44.2 | 45.2 |
| *Alcaligenes faecalis* | P | Negative | 44.39 | 40.8 | 0.714 | 0.6238 | 56.81 | 57.5 |
| *Aliivibrio wodanis* | P | Negative | 45.7 | 41.07 | 0.273 | 0.1973 | 38.3 | 43.25 |
| *Alteromonas macleodii* ATCC 27126 | NP | Negative | 53.15 | 42.82 | 0.411 | 0.2791 | 44.7 | 48.16 |
| *Anaerostipes hadrus* DSM 3319 | NP | Positive | 42.424 | 41.37 | 0.257 | 0.209 | 37.1 | 44.13 |
| *Anaplasma marginale* str. Florida | P | Negative | 55.68 | 56.88 | 0.52 | 0.5459 | 49.8 | 51.49 |
| *Anoxybacillus gonensis* | NP | Positive | 47.027 | 42.2 | 0.443 | 0.5079 | 41.7 | 48.08 |
| *Arcobacter butzleri* RM4018 | P | Negative | 33.144 | 30.85 | 0.0568 | 0.01974 | 27 | 31.31 |
| *Arthrobacter* sp. ATCC 21022 | NP | Positive | 41.9 | 35.98 | 0.776 | 0.7443 | 64.5 | 66.18 |
| *Bacillus anthracis* str. Ames | P | Positive | 43.808 | 36.21 | 0.249 | 0.1347 | 35.4 | 40.48 |
| *Bacillus mycoides* | NP | Positive | 44.08 | 38.08 | 0.249 | 0.1214 | 35.2 | 32.92 |
| *Bacteroides cellulosilyticus* | P | Negative | 50.17 | 45.97 | 0.435 | 0.61 | 42.7 | 50.45 |
| *Bartonella bacilliformis* KC583 | P | Negative | 44.84 | 38.92 | 0.279 | 0.2074 | 38.2 | 43.23 |
| *Bifidobacterium adolescentis* ATCC 15703 | NP | Positive | 40.339 | 33.14 | 0.765 | 0.7975 | 59.2 | 63.58 |
| *Blautia obeum* | NP | Positive | 45.546 | 42.21 | 0.36 | 0.2439 | 41.2 | 45.97 |
| *Bordetella bronchiseptica* 253 | P | Negative | 33.148 | 29.98 | 0.9329 | 0.9196 | 68.1 | 66.67 |
| *Borrelia burgdorferi* B31 | P | Negative | 40.56 | 36.28 | 0.181 | 0.1946 | 28.18 | 37.83 |
| *Brevibacillus brevis* NBRC 100599 | NP | Positive | 54.223 | 46.54 | 0.511 | 0.3824 | 47.3 | 50.39 |
| *Brucella melitensis* bv. 1 str. 16M | P | Negative | 44.698 | 54.33 | 0.724 | 0.7538 | 57.24 | 58.74 |
| *Buchnera aphidicola* str. APS (*Acyrtosiphon pisum*) | NP | Negative | 37.33 | 36.7 | 0.131 | 0.1479 | 26.4 | 33.94 |
| *Burkholderia gladioli* | P | Negative | 34.048 | 29.7 | 0.9136 | 0.9593 | 68 | 70.78 |
| *Burkholderia ubonensis* MSMB22 | NP | Negative | 33.516 | 26.99 | 0.92 | 0.9447 | 67.31 | 67.42 |
| *Butyrivibrio proteoclasticus* B316 | NP | Positive | 46.349 | 36.02 | 0.29 | 0.1473 | 40 | 45.01 |
| *Caldicellulosiruptor bescii* DSM 6725 | NP | Positive | 44.89 | 49.39 | 0.238 | 0.2821 | 35.22 | 40.07 |
| *Capnocytophaga ochracea* DSM 7271 | P | Negative | 48.47 | 44.39 | 0.36 | 0.2059 | 39.6 | 37.26 |
| *Caulobacter crescentus* CB15 | NP | Negative | 35.29 | 29.63 | 0.883 | 0.8675 | 67.2 | 67.58 |
| *Chania multitudinisentens* RB-25 | NP | Negative | 48.29 | 42.16 | 0.616 | 0.5896 | 50.9 | 54.72 |
| *Chlamydophila pneumoniae* CWL029 | P | Negative | 50.47 | 61 | 0.326 | 0.3846 | 40.6 | 41.29 |
| *Chromobacterium subtsugae* | NP | Negative | 33.41 | 30.35 | 0.916 | 0.9539 | 64.8 | 67.41 |
| *Chronobacter sakazakii* | P | Negative | 42.83 | 39.1 | 0.744 | 0.6861 | 56.9 | 58.16 |
| *Citrobacter amalonaticus* | P | Negative | 46.56 | 40.25 | 0.672 | 0.653 | 53.21 | 56.94 |
| *Clavibacter michiganensis* subsp. sepedonicus | NP | Positive | 30.78 | 27.21 | 0.932 | 0.9576 | 72.4 | 71.58 |
| *Clostridium bolteae* 90A9 | P | Positive | 48.257 | 47.82 | 0.6061 | 0.5505 | 49.6 | 52.21 |
| *Clostridium butyricum* | NP | Positive | 36 | 33.83 | 0.1 | 0.09884 | 28.6 | 35.8 |
| *Comamonas testosteroni* TK102 | P | Negative | 39.89 | 34.91 | 0.8 | 0.8059 | 61.9 | 63.74 |
| *Corynebacterium diphtheriae* | P | Positive | 48.411 | 49.9 | 0.538 | 0.405 | 53.6 | 52.91 |
| *Corynebacterium glutamicum* ATCC 13032 | NP | Positive | 47.47 | 46.74 | 0.56 | 0.3267 | 53.8 | 59.52 |
| *Coxiella burnetii* RSA 493 | P | Negative | 52.24 | 53.76 | 0.49 | 0.5665 | 42.64 | 49.92 |
| *Cupriavidus metallidurans* CH34 | NP | Negative | 40.37 | 30.27 | 0.81 | 0.8745 | 63.52 | 64.99 |
| *Cutibacterium avidum* 44067 | P | Positive | 40.52 | 36.31 | 0.764 | 0.7449 | 63.5 | 64.51 |
| *Deinococcus radiodurans* R1 | NP | Positive | 37.44 | 28.77 | 0.8732 | 0.9393 | 66.7 | 66.76 |
| *Delftia acidovorans* SPH-1 | P | Negative | 34.39 | 30.84 | 0.88 | 0.9188 | 66.5 | 70.6 |
| *Desulfovibrio vulgaris* str. Hildenborough | P | Negative | 42.051 | 36.38 | 0.763 | 0.7705 | 63.24 | 63.04 |
| *Edwardsiella ictaluri* 93-146 | P | Negative | 43.07 | 38.45 | 0.73 | 0.6878 | 57.4 | 58.31 |
| *Eikenella corrodens* ATCC 23834 | P | Negative | 41.16 | 38.32 | 0.714 | 0.6146 | 55.7 | 53.88 |
| *Eisenbergiella tayi* | NP | Negative | 48.215 | 46.42 | 0.586 | 0.5222 | 46.8 | 51.42 |
| *Ensifer adhaerens* | NP | Negative | 40.38 | 29.21 | 0.815 | 0.8272 | 62.2 | 67.06 |
| *Enterobacter (Klebsiella) aerogenes* KCTC 2190 | P | Negative | 43.77 | 37.36 | 0.7223 | 0.6441 | 54.8 | 57.2 |
| *Enterococcus avium* ATCC 14025 | P | Positive | 50.33 | 38.28 | 0.372 | 0.1786 | 39.1 | 41.24 |
| *Escherichia coli* IAI39 | NP | Negative | 47.96 | 44.85 | 0.606 | 0.5091 | 50.6 | 53.82 |
| *Flavobacterium hydatis* | P | Negative | 43.47 | 39.39 | 0.2107 | 0.1213 | 32.7 | 54.97 |
| *Francisella philomiragia* subsp. *philomiragia* ATCC 25017 | P | Negative | 41.0375 | 33.83 | 0.15 | 0.08808 | 32.59 | 39.15 |
| *Fusobacterium nucleatum* | P | Negative | 33.153 | 30.57 | 0.08 | 0.02721 | 27 | 32.04 |
| *Gallibacterium anatis* UMN179 | P | Negative | 45.371 | 40 | 0.407 | 0.3452 | 39.89 | 42.96 |
| *Gardnerella vaginalis* 409-05 | P | Gram Variable | 44.71 | 43.91 | 0.281 | 0.2315 | 42 | 47.08 |
| *Geobacillus subterraneus* | NP | Positive | 42.583 | 43.67 | 0.751 | 0.7788 | 52.2 | 57.58 |
| *Geobacter sulfurreducens* PCA | NP | Negative | 42.76 | 33.86 | 0.791 | 0.8571 | 60.9 | 62.15 |
| *Gluconobacter oxydans* 621H | NP | Negative | 42.58 | 38.42 | 0.751 | 0.7771 | 60.84 | 65.17 |
| *Granulibacter bethesdensis* CGDNIH1 | P | Negative | 46.099 | 37.13 | 0.707 | 0.8012 | 59.1 | 66.06 |
| *Haemophilus influenzae* Rd KW20 | P | Negative | 43.909 | 44.54 | 0.329 | 0.244 | 38.2 | 41 |
| *Halomonas boliviensis* LC1 | NP | Negative | 49.93 | 50.66 | 0.644 | 0.5198 | 54.6 | 55.53 |
| *Helicobacter pylori* 26695 | P | Negative | 47.028 | 44.64 | 0.505 | 0.4167 | 38.9 | 43.78 |
| *Ketogulonicigenium vulgare* WSH-001 | NP | Negative | 40.641 | 39.63 | 0.816 | 0.7981 | 61.73 | 64.03 |
| *Klebsiella oxytoca* | P | Negative | 44.152 | 39.72 | 0.728 | 0.6802 | 55.2 | 58.42 |
| *Kocuria kristinae* | P | Positive | 29.198 | 24.56 | 0.947 | 0.9577 | 71.8 | 70.75 |
| *Lactobacillus amylovorus* | NP | Positive | 44.77 | 44.07 | 0.262 | 0.1527 | 38.08 | 41.21 |
| *Lactobacillus crispatus* ST1 | P | Positive | 44.95 | 36.61 | 0.273 | 0.1238 | 36.9 | 41.67 |
| *Lactococcus garvieae* Lg2 | P | Positive | 47.9 | 34.19 | 0.311 | 0.161 | 38.8 | 43.29 |
| *Lactococcus lactis* subsp. *lactis* II1403 | NP | Positive | 43.739 | 33.67 | 0.24 | 0.1161 | 35.3 | 41.95 |
| *Methylobacterium aquaticum* | NP | Negative | 33.251 | 26.96 | 0.921 | 0.9819 | 70.9 | 73.81 |
| *Microbacterium foliorum* | NP | Positive | 34.69 | 30.91 | 0.87 | 0.8455 | 67.9 | 67.8 |
| *Micrococcus luteus* NCTC 2665 | P | Positive | 30.032 | 26.34 | 0.945 | 0.9731 | 73 | 73.85 |
| *Moraxella catarrhalis* BBH18 | P | Negative | 47.72 | 44.35 | 0.403 | 0.3443 | 41.7 | 44.92 |
| *Morganella morganii* subsp. *morganii* KT | P | Negative | 43.39 | 38.92 | 0.653 | 0.6038 | 51.1 | 54.49 |
| *Mycobacterium abscessus* | P | Positive | 42.411 | 30.91 | 0.795 | 0.8455 | 64.1 | 67.87 |
| *Neisseria gonorrhoeae* FA 1090 | P | Negative | 44.08 | 43.97 | 0.69 | 0.6 | 52.7 | 52.67 |
| *Neorhizobium galegae* bv. *orientalis* str. HAMBI 540 | NP | Negative | 41.17 | 32.05 | 0.81 | 0.7837 | 61.25 | 65.83 |
| *Obesumbacterium proteus* | NP | Negative | 49.55 | 40.26 | 0.55 | 0.467 | 49.05 | 52.63 |
| *Ochrobactrum anthropi* ATCC 49188 | NP | Negative | 46.8 | 36.7 | 0.675 | 0.6106 | 56.15 | 61.66 |
| *Oenococcus oeni* PSU-1 | NP | Positive | 49.59 | 48.33 | 0.36 | 0.3065 | 37.9 | 44.23 |
| *Orientia tsutsugamushi* str. Boryong | P | Negative | 42.74 | 36.9 | 0.18 | 0.1351 | 30.5 | 35.02 |
| *Pantoea ananatis* LMG 20103 | NP | Negative | 47.16 | 42.46 | 0.672 | 0.5794 | 53.7 | 55.24 |
| *Phaeobacter gallaeciensis* DSM 26640 | NP | Negative | 45.35 | 41.71 | 0.738 | 0.7275 | 59.42 | 63.33 |
| *Photobacterium kishitanii* | NP | Negative | 44.94 | 39.99 | 0.309 | 0.2 | 38.8 | 45.09 |
| *Photorhabdus temperata* subsp. *thracensis* | NP | Negative | 50.35 | 44.89 | 0.452 | 0.4028 | 44.1 | 48.92 |
| *Piscirickettsia salmonis* LF-89 = ATCC VR-1361 | P | Negative | 48 | 45.74 | 0.366 | 0.4731 | 39.62 | 43.83 |
| *Pluralibacter gergoviae* | P | Negative | 39.67 | 36.02 | 0.807 | 0.7534 | 59 | 59.2 |
| *Polynucleobacter asymbioticus* QLW-P1DMWA-1 | NP | Negative | 51.531 | 41.18 | 0.4 | 0.2448 | 44.8 | 47.05 |
| *Porphyromonas gingivalis* ATCC 33277 | P | Negative | 52.9 | 51.39 | 0.543 | 0.6288 | 48.4 | 52.26 |
| *Prevotella melaninogenica* ATCC 25845 | P | Negative | 48.031 | 41.73 | 0.29 | 0.3109 | 40.99 | 47.93 |
| *Prevotella ruminicola* 23 | NP | Negative | 46.06 | 41.02 | 0.505 | 0.4396 | 47.7 | 51.75 |
| *Prochlorococcus marinus* str. AS9601 | NP | Negative | 40.69 | 37.8 | 0.173 | 0.2081 | 31.3 | 39.78 |
| *Propionibacterium acnes* KPA171202 | NP | Positive | 48.063 | 45.69 | 0.67 | 0.5992 | 60 | 60.85 |
| *Proteus mirabilis* HI4320 | P | Negative | 46.533 | 40.77 | 0.344 | 0.3184 | 38.88 | 45.24 |
| *Providencia stuartii* MRSN 2154 | P | Negative | 49.64 | 45.56 | 0.392 | 0.3684 | 41.3 | 47.98 |
| *Pseudoalteromonas luteoviolacea* | NP | Negative | 51.78 | 47.2 | 0.372 | 0.3288 | 42 | 46.06 |
| *Ralstonia pickettii* 12J | P | Negative | 38.84 | 31.7 | 0.831 | 0.8312 | 63.62 | 64.17 |
| *Ralstonia solanacearum* GMI1000 | NP | Negative | 34.87 | 28.7 | 0.896 | 0.9603 | 66.96 | 70.33 |
| *Rhizobium etli* CFN 42 | NP | Negative | 42.22 | 33.46 | 0.802 | 0.7994 | 61.05 | 65.28 |
| *Rhodanobacter thiooxydans* | NP | Negative | 33.406 | 29.32 | 0.898 | 0.9163 | 67.2 | 67.85 |
| *Rhodobacter sphaeroides* 2.4.1 | NP | Negative | 34.94 | 29.33 | 0.914 | 0.9228 | 68.78 | 70.95 |
| *Rhodococcus aetherivorans* | NP | Positive | 33.051 | 29.55 | 0.913 | 0.9336 | 70.44 | 69.67 |
| *Rhodococcus equi* 103S | P | Positive | 34.749 | 30.2 | 0.878 | 0.8315 | 68.8 | 71.68 |
| *Rhodospirillum rubrum* ATCC 11170 | NP | Negative | 37.7 | 35.01 | 0.88 | 0.8929 | 65.33 | 69.87 |
| *Rickettsia conorii* str. Malish 7 | P | Negative | 44.194 | 42.27 | 0.23 | 0.2379 | 32.4 | 38.04 |
| *Riemerella anatipestifer* ATCC 11845 = DSM 15868 | NP | Negative | 45.19 | 42.56 | 0.257 | 0.1646 | 35 | 36.49 |
| *Rothia dentocariosa* | P | Positive | 48.9 | 40.33 | 0.58 | 0.496 | 53.8 | 57.84 |
| *Rothia dentocariosa* ATCC 17931 | NP | Positive | 49.34 | 47.28 | 0.575 | 0.448 | 53.7 | 55.45 |
| *Salinibacter ruber* DSM 13855 | NP | Negative | 38.68 | 31.59 | 0.864 | 0.9605 | 66.12 | 67.12 |
| *Salinispora tropica* CNB-440 | NP | Positive | 36.56 | 35.49 | 0.85 | 0.7911 | 69.5 | 66.49 |
| *Salmonella enterica* subsp. enterica serovar Typhi str. CT18 | P | Negative | 47.73 | 41.55 | 0.646 | 0.6818 | 51.88 | 57.64 |
| *Selenomonas noxia* ATCC 43541 | NP | Negative | 44.83 | 38 | 0.69 | 0.7037 | 55.8 | 60.93 |
| *Serratia fonticola* | NP | Negative | 46.17 | 37.3 | 0.69 | 0.6712 | 53.8 | 57.84 |
| *Serratia rubidaea* | P | Negative | 38.52 | 36.06 | 0.836 | 0.7281 | 59.25 | 58.4 |
| *Shewanella baltica* OS678 | NP | Negative | 50.59 | 45.79 | 0.537 | 0.4018 | 46.28 | 50.76 |
| *Shigella dysenteriae* Sd197 | NP | Negative | 48.9 | 44.82 | 0.613 | 0.5273 | 50.92 | 54.17 |
| *Shigella flexneri* 2a str. 301 | P | Negative | 48.64 | 44.97 | 0.603 | 0.5291 | 50.67 | 54.17 |
| *Sinorhizobium fredii* NGR234 | NP | Negative | 41.64 | 33.64 | 0.8167 | 0.8261 | 62.4 | 66.27 |
| *Staphylococcus aureus* | P | Positive | 41.019 | 34.09 | 0.207 | 0.07407 | 32.7 | 38.53 |
| *Staphylococcus capitis* subsp. capitis | NP | Positive | 41.48 | 37.79 | 0.192 | 0.1179 | 32.94 | 39.98 |
| *Stenotrophomonas maltophilia* | P | Negative | 32.627 | 26.77 | 0.88 | 0.8976 | 66.4 | 68.12 |
| *Streptococcus agalactiae* 2603V/R | NP | Positive | 44.496 | 36.22 | 0.22 | 0.1273 | 35.6 | 40.05 |
| *Streptococcus pneumoniae* R6 | P | Positive | 47.61 | 38.53 | 0.314 | 0.193 | 39.7 | 44.52 |
| *Streptomyces lydicus* | NP | Positive | 31.6 | 28.92 | 0.928 | 0.9067 | 72.05 | 71.13 |
| *Thioalkalivibrio versutus* | NP | Negative | 36.13 | 33.78 | 0.874 | 0.8987 | 66.2 | 64.94 |
| *Treponema denticola* ATCC 35405 | NP | Negative | 49.47 | 42.35 | 0.37 | 0.2383 | 37.9 | 39.8 |
| *Tropheryma whipplei* str. Twist | P | Positive | 54.87 | 52.62 | 0.429 | 0.4834 | 46.3 | 51.47 |
| *Vibrio alginolyticus* NBRC 15630 = ATCC 17749 | P | Negative | 50.85 | 45.75 | 0.401 | 0.2763 | 44.7 | 48.02 |
| *Weissella cibaria* | NP | Positive | 45.69 | 37.12 | 0.414 | 0.308 | 44.9 | 48.74 |
| *Wolbachia endosymbiont of Drosophila melanogaster* | NP | Negative | 46.88 | 41.99 | 0.245 | 0.1648 | 35.2 | 38.4 |
| *Xanthomonas campestris* pv. campestris str. ATCC 33913 | NP | Negative | 36.64 | 30.17 | 0.853 | 0.8745 | 65.1 | 66.42 |
| *Xenorhabdus bovienii* SS-2004 | NP | Negative | 51.25 | 48.23 | 0.475 | 0.4486 | 45 | 49.91 |
| *Xylella fastidiosa* 9a5c | NP | Negative | 48.926 | 42.99 | 0.556 | 0.4735 | 52.64 | 54.45 |
| *Yersinia aldovae* | NP | Negative | 50.561 | 48.11 | 0.541 | 0.4928 | 47.7 | 51.91 |
| *Yersinia pestis* CO92 | P | Negative | 51.021 | 45.27 | 0.54 | 0.5117 | 47.61 | 52.52 |

After analysing the codon usage data of *ftsZ* from the 142 species we found that *Kocuria kristinae*, which is a pathogenic, Gram-positive bacteria exhibits the lowest Nc value of 24.65 among all the organisms. On the other hand, *Chlamydophila pneumoniae* CWL 029, a pathogenic, Gram-negative bacteria exhibited the highest Nc value of 61. A higher Nc value indicates poor codon bias of the gene [36]. Analysing the mean genomic Nc value of all the organisms studied, it was observed that the lowest mean genomic Nc value (29.198) is depicted by the organism *Kocuria kristinae*, a pathogenic, Gram-positive bacteria whereas the maximum mean genomic Nc value (55.68) is depicted by a pathogenic, Gram-negative bacteria called *Anaplasma marginale* str. Florida. Our observations primarily suggest that the degree of codon bias in the pathogenic organisms span a wider range.

Following the trend in Nc values, we clearly observed that the mean genomic Nc in majority of the organisms is higher than the genic Nc of *ftsZ*. This suggests that the *ftsZ* gene is subjected to greater codon bias in comparison to the genomic Nc. But in case of nine organisms, exceptions were evident. These organisms include *Haemophilus influenzae* Rd KW20, *Halomonas boliviensis* LC1, *Geobacillus subterraneus*, *Anaplasma marginale* str. Florida, *Corynebacterium diphtheria*, *Coxiella burnetii* RSA 493, *Caldicellulosiruptor bescii* DSM 6725, *Brucella melitensis* bv. 1 str. 16M and *Chlamydophila pneumoniae* CWL029. Most of these organisms are Gram negative and pathogenic in nature.

In case of GC3 content, *Arcobacter butzleri* RM 4018, a pathogenic Gram negative strain depicted the lowest GC3 value for *ftsZ* gene (0.01974). The maximum GC3 content for *ftsZ* was shown by *Methylobacterium aquaticum* (0.9819), a non-pathogenic Gram negative bacteria.

Statistical analysis demonstrated a significant positive correlation between the mean genomic Nc and *ftsZ* genic Nc by Spearman’s Rank correlation (*ρ*=0.863, p<<0.01). We have also detected a significant negative correlation between the Nc and GC3 of the *ftsZ* gene ((*ρ*=-0.491, p<<0.01) by Spearman’s rank correlation.

The study of the relation between Nc and GC3 is an important analytical tool for examining codon bias. So, to better understand the codon usage bias profile of the *ftsZ* genes an Nc-plot was constructed. Analysis of the Nc-plot shows that the *ftsZ* genes of three pathogenic organisms— *Anaplasma marginale* str. Florida, *Brucella melitensis* bv. 1 str. 16M and *Chlamydophila pneumoniae* CWL029 occupy distinct positions on the Nc-plot (Fig 1). The common features shared by these three organisms are that they are Gram negative and pathogenic in nature. The bacteria *Anaplasma marginale* is a member of the order Rickettsiales. It is a small, obligate intracellular bacteria that typically have short genomes due to reductive evolution and survive as endosymbionts. It is also responsible for an infectious, noncontagious disease called bovine anaplasmosis in cattle and other ruminants [37]. The other organism *Brucella melitensis* is responsible for brucellosis, a common health hazard in people living in close vicinity of cattle [38]. The third organism called *Chlamydophila pneumoniae* represents an intracellular pathogen instigating different acute and chronic infections and has been found to be associated with chronic neurological disorders such as Alzheimer’s disease and multiple sclerosis. Infection by *C. pneumoniae* which is a common cause of human respiratory disease [39] has also been suspected to cause chronic fatigue syndrome and the linked syndrome polymyalgia rheumatic in some patients [40].

**Fig 1:**
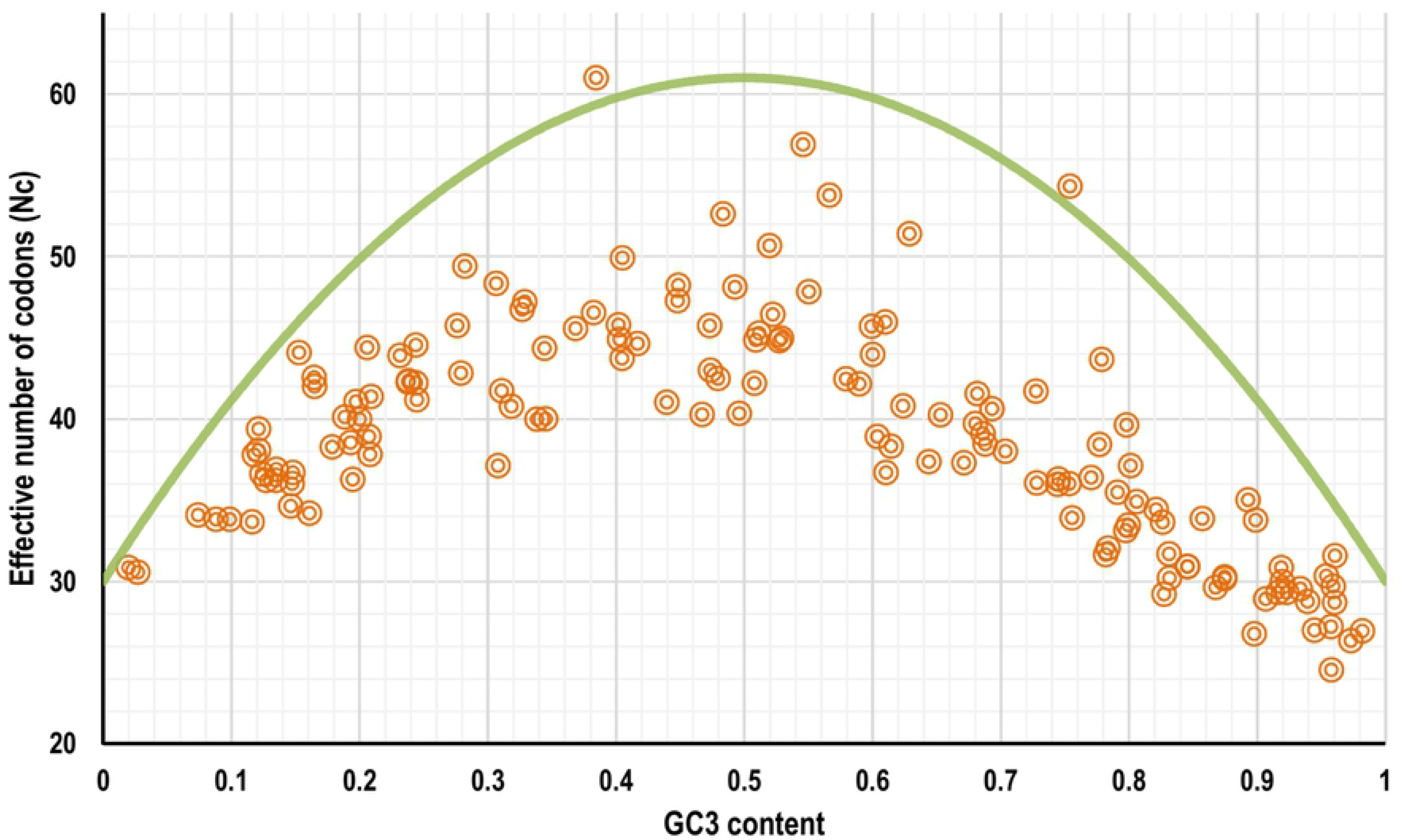
An Nc-plot depicting the correlation between Nc and GC3 of the 143 *ftsZ* genes selected from 143 different bacteria. The continuous curve depicts the null hypothesis that the GC bias at the synonymous site is solely due to mutation but not selection.

In order to study the compositional divergence of the gene sequences coding for FtsZ protein in the selected organisms with respect to their whole genome, the difference between the mean genomic GC content with the GC content of *ftsZ* gene, mean genomic Nc with Nc of *ftsZ* and the difference between average whole genome GC3 content with that of the *ftsZ* coding sequence was analysed.

### Difference between mean genomic GC and *ftsZ* GC

The guanine-cytosine (GC) composition of bacterial genomes is a very important taxonomic marker from the genomics perspective. The GC content of a genome as well as that of a gene have been reported to be a significant genomic indicator for comparison between covalently closed circular plasmid DNA and chromosomes [41], and for distinguishing between vertically and horizontally transferred genes [42]. In our study we found that, out of the 143 organisms, the *ftsZ* CDS of 49 organisms depicted greater than 10% GC skew in comparison to their genomic GC content. Among these organisms, *Coxiella burnetii* RSA 493, *Rickettsia conorii* str. Malish 7, *Staphylococcus aureus*, *Bacteroides cellulosilyticus*, *Fusobacterium nucleatum*, *Lactococcus lactis* subsp. lactis Il1403, *Anaerostipes hadrus* DSM 3319 and *Acinetobacter johnsonii* XBB1 demonstrates 15% greater usage of guanine and cytosine residues in their *ftsZ* CDS in comparison to the whole genome GC content. In comparison to the genomic GC content, a relatively greater usage of guanine and cytosine residues (more than 20% to 68%) was observed in the *ftsZ* CDS of *Francisella philomiragia* subsp. philomiragia ATCC 25017, *Staphylococcus capitis* subsp. capitis, *Clostridium butyricum*, *Prochlorococcus marinus* str. AS9601, *Buchnera aphidicola* str. APS, *Borrelia burgdorferi* B31 and *Flavobacterium hydatis*. The GC content of *ftsZ* CDS in comparison to the genomic GC of *Flavobacterium hydatis* was an extraordinarily 68% greater. On the other hand, the GC content of *ftsZ* CDS in comparison to the genomic GC content of organisms like *Bacillus mycoides*, *Capnocytophaga ochracea* DSM 7271, *Salinispora tropica* CNB-440, *Eikenella corrodens* ATCC 23834 and *Bordetella bronchiseptica* 253 was found to be 2% to 6% lower. A Mann-Whitney U test was conducted to statistically validate the difference between the genomic GC content and the *ftsZ* genic GC content of the 143 species. The results suggest that the genomic GC content and *ftsZ* GC content differs significantly (*U*=8536.50, *p*=0.016). All of the above findings clearly suggest that the nucleotide composition of the gene coding for FtsZ protein in a large number of species deviates significantly from their genomic GC content. The deviation of GC content of a coding sequence or a patch of nucleotides from the genomic GC content is a possible pointer towards horizontal gene transfer or HGT [43], and our analysis using Mann-Whitney U test also points in that direction.

### Difference between mean genomic Nc and *ftsZ* Nc

Out of the 143 organisms, about 93% (134 species) demonstrated relatively biased codon usage configuration in terms of Nc value. Of these 143 organisms, 21 species *viz.*, *Lactococcus garvieae* Lg2, *Ensifer adhaerens*, *Mycobacterium abscessus*, *Aerococcus viridans*, *Cupriavidus metallidurans* CH34, *Enterococcus avium* ATCC 14025, *Deinococcus radiodurans* R1, *Lactococcus lactis* subsp. lactis Il1403, *Butyrivibrio proteoclasticus* B316, *Neorhizobium galegae* bv. orientalis str. HAMBI 540, *Ochrobactrum anthropi* ATCC 49188, *Geobacter sulfurreducens* PCA, *Rhizobium etli* CFN 42, *Polynucleobacter asymbioticus* QLW-P1DMWA-1, *Burkholderia ubonensis* MSMB22, *Granulibacter bethesdensis* CGDNIH1, *Alteromonas macleodii* ATCC 27126, *Sinorhizobium fredii* NGR234, *Serratia fonticola* and *Streptococcus pneumoniae* R6 demonstrated Nc values of *ftsZ* coding sequences that are 20% or less than their mean genomic Nc values. This is suggestive of a significant codon bias existing within the *ftsZ* CDS. On the other hand, the *ftsZ* CDS of two Gram negative and pathogenic species *Chlamydophila pneumoniae* CWL029 and *Brucella melitensis* bv. 1 str. 16M were found to display Nc values twenty units greater than their mean genomic Nc score.

### Difference between mean genomic GC3 and genic *ftsZ* GC3

Out of the 143 organisms, the *ftsZ* CDS of organisms like *Fusobacterium nucleatum*, *Arcobacter butzleri* RM4018, *Staphylococcus aureus*, *Aerococcus viridans*, *Lactobacillus crispatus* ST1, *Enterococcus avium* ATCC 14025, *Lactococcus lactis* subsp. lactis Il1403, *Bacillus mycoides*, *Butyrivibrio proteoclasticus* B316 had GC3 content which was substantially less (upto 66% lesser) than the mean genomic GC3 content. Barring *Lactococcus lactis* subsp. lactis Il1403, *Bacillus mycoides* and *Butyrivibrio proteoclasticus* B316, the remaining organisms are pathogenic in nature. This is an interesting observation which shows that the *ftsZ* ORFs of these pathogenic bacteria are structured without significant bias towards G and C ending codons. Organisms like *Prochlorococcus marinus* str. AS9601, *Piscirickettsia salmonis* LF-89 and *Bacteroides cellulosilyticus* on the other hand, had significantly greater GC3 (20%, 29% and 40% respectively) in their *ftsZ* CDS compared to their genomic GC3 content.

### Analysis of codon usage to detect ‘core’ set of codons used in structuring of *ftsZ*

The individual usage frequency of the 61 sense codons from the 143 organisms were calculated. Out of the 61 sense codons, the two non-degenerate codons coding for methionine and tryptophan were eliminated. For the remaining 18 amino acids, the 59 codons were grouped in to their degenerate classes of 2, 3, 4 and 6 codons. This analysis was performed to find out if there exists a preferred set of ‘core’ codons for each of the amino acids used in structuring of the *ftsZ* CDS. A Kruskal-Wallis one way analysis of variance on ranks was carried out for the amino acids coded by 3, 4 and 6 codons, whereas Mann-Whitney Rank Sum test was used to test the codon preference in the two codon family amino acids. The results established the fact that, out of the 18 amino acids, the codons of three amino acids namely aspartic acid, histidine and alanine are randomly utilised on a global scale for structuring the *ftsZ* CDS. On the other hand, the codons for the remaining 15 amino acids show a non-random utilization pattern. These amino acids include cysteine, glutamine, phenylalanine, glycine, isoleucine, lysine, leucine, asparagine, proline, glutamine, arginine, serine, threonine, valine and tyrosine. Table 2 contains the Mann-Whitney U statistic and the H-value with degrees of freedom for the Kruskal-Wallis one way analysis of variance on ranks with their corresponding *p-*value obtained from the tests. Our analysis using both the above mentioned robust inferential statistical tools suggest that for all the 18 amino acids (except aspartic acid, histidine and alanine) the differences in the median values among the codon groups are greater than would be expected by chance and hence there is a statistically significant difference at *p=*<0.001 level. This is an important finding suggesting the antiquity and conservation of a preferred set of codons in structuring of a vital gene such as the *ftsZ* gene.

**Table 2:** Mann-Whitney U statistic and the H-value with degrees of freedom for the Kruskal-Wallis one way ANOVA on ranks on codon usage to detect ‘core’ set of codons used in structuring of *ftsZ*.

| Sl. No. | Amino acid | Degenerate codon family | Mann-Whitney U statistic | H-value with degrees of freedom (df) for the Kruskal-Wallis one way analysis of variance on ranks | p-value |
| --- | --- | --- | --- | --- | --- |
| 1 | Cys | 2 | 8029.50 | - | <0.001 |
| 2 | Glu | 2 | 5622.00 | - | <0.001 |
| 3 | Phe | 2 | 8375.50 | - | 0.008 |
| 4 | Lys | 2 | 7552.00 | - | <0.001 |
| 5 | Asn | 2 | 6426.00 | - | <0.001 |
| 6 | Gln | 2 | 6384.00 | - | <0.001 |
| 7 | His | 2 | 9211.00 | - | 0.145 |
| 8 | Asp | 2 | 9565.50 | - | 0.346 |
| 9 | Tyr | 2 | 8696.50 | - | 0.001 |
| 10 | Ile | 3 | - | 229.853, *df*=2 | <0.001 |
| 11 | Gly | 4 | - | 273.266, *df*=3 | <0.001 |
| 12 | Thr | 4 | - | 70.997, *df*=3 | <0.001 |
| 13 | Val | 4 | - | 49.583, *df*=3 | <0.001 |
| 14 | Ala | 4 | - | 3.777, *df*=3 | 0.287 |
| 15 | Pro | 4 | - | 69.438, *df*=3 | <0.001 |
| 16 | Leu | 6 | - | 144.051, *df*=5 | <0.001 |
| 17 | Arg | 6 | - | 406.291, *df*=5 | <0.001 |
| 18 | Ser | 6 | - | 76.847, *df*=5 | <0.001 |

### Two factor ANOVA on codon usage of *ftsZ* CDS to study the relationship of the frequency of the individual 61 sense codons and their interrelation with lifestyle and Gram nature of bacteria

Sixty one separate variance analysis tests called two factor (or two way) ANOVA was performed to find out how the two major factors namely lifestyle (pathogenic or non-pathogenic), Gram nature and interaction of these two factors influence the coding composition of the *ftsZ* CDS in the selected organisms at *p*<0.01 level of significance. A critical analysis of the results show that the compositional bias of eight codons coding for six amino acids is influenced mostly by the Gram nature of the organisms and in some instances by the interaction of lifestyle and Gram nature. In our study, we find that the compositional bias of the codons AUG (methionine), UCA (serine), UAU (tyrosine) and UAC (tyrosine) is influenced solely by the Gram nature of the organism. On the other hand, the compositional frequency of the codons GGA (glycine), CUU (leucine), CUG (leucine) and ACA (threonine) is influenced by the interaction between the Gram nature of the organism and their lifestyle preference of being either pathogenic or non-pathogenic. The two way ANOVA results suggest that the codon organization of the *ftsZ* CDS is determined largely by the Gram nature and pathogenic/non-pathogenic nature of the organisms, and it is a non-randomly constituted sequence in terms of codon deployment.

### Utilization of two factor ANOVA on *ftsZ* CDS to study the frequency of the individual 20 amino acids and their interrelation with lifestyle and Gram nature

To further comprehend the codon deployment pattern of the *ftsZ* CDS, a two way ANOVA was carried out by grouping the different codons according to the amino acids they code (for example alanine is coded by four codons and these four codons are clubbed into a single category to estimate the total frequency of alanine residues present in the CDS). Twenty discrete two way ANOVA analysis was carried out to find if the two factors namely lifestyle, Gram nature and interaction of these two factors influence the amino acid composition of the *ftsZ* CDS in the selected organisms (at *p*<0.01 level of significance) or, is the amino acid composition random in nature. All the post-hoc pairwise multiple comparison in the analysis was performed using the Holm-Sidak method of pairwise multiple comparison [44, 45]. The results show that the compositional frequency of the amino acids glutamic acid, phenylalanine, leucine, valine, glutamine, threonine and tryptophan is influenced neither by the lifestyle nor the Gram nature of the organism. But, the frequency of the amino acids like aspartic acid, histidine, glycine, methionine, cysteine and tyrosine is influenced by the Gram nature of the organism (*p*<0.01 level). This shows that the compositional frequency of at least one amino acid from the four different chemical classes of amino acids is directly associated with the Gram nature of the bacteria. Another interesting observation is that the two sulphur containing amino acids methionine and cysteine are both involved in inducing compositional variability based on the wall nature of the bacterium. The hydroxymethyl side chain containing polar amino acid serine was found to be unique in the sense that a two factor ANOVA on composition frequency of serine detected that it is influenced both by the Gram nature and lifestyle of the organism. No amino acid other than serine was found to be influenced by the lifestyle of the organism. Thus, serine appears to be the only amino acid in the FtsZ protein which acts as a marker of the lifestyle of the bacterial species considered in this study. In case of the compositional frequency of the remaining amino acids like alanine, isoleucine, proline, lysine, arginine and asparagine the effect of lifestyle was found to rest on the Gram nature of the organisms at *p*<0.01 level.

### Identity and cluster based analysis of *ftsZ* CDS

The sequence identity of the 143 FtsZ proteins were determined using Clustal Omega [32]. We observed that the identity of the FtsZ proteins fluctuated tremendously among the different bacterial species. The identity was found to range from 13% to 93% among the organisms selected for this study. The FtsZ protein of organisms like *Pluralibacter gergoviae, Chronobacter sakazakii, Shigella flexneri* 2a str. 301*, Salmonella enterica* subsp. enterica serovar Typhi str. CT18*, Klebsiella oxytoca, Citrobacter amalonaticus, Enterobacter aerogenes* KCTC 2190*, Escherichia coli* IAI39*, Shigella dysenteriae* Sd197*, Edwardsiella ictaluri* 93-146*, Obesumbacterium proteus, Yersinia aldovae, Yersinia pestis* CO92*, Serratia rubidaea, Pantoea ananatis* LMG 20103*, Chania multitudinisentens* RB-25*, Serratia fonticola, Photorhabdus temperata* subsp. thracensis*, Xenorhabdus bovienii* SS-2004*, Morganella morganii* subsp. morganii KT*, Proteus mirabilis* HI4320 and *Providencia stuartii* MRSN 2154 was found to share greater than 90% identity. On the other hand organisms like *Chromobacterium subtsugae*, *Helicobacter pylori* 26695, *Arcobacter butzleri* RM4018, *Ralstonia solanacearum* GMI1000 and *Fusobacterium nucleatum* was found to share less than 15% identity in their FtsZ protein sequences.

The 143 *ftsZ* CDS were subjected to clustering using CD-HIT with a 50% similarity threshold. A tabular account of the 17 clusters generated using CD-HIT along with the number of representative sequences for each cluster is given in Table 3. From the data given in Table 3, it is quite evident that the majority of the sequences are grouped together in the first two clusters which contains 43% of the total *ftsZ* CDS (41 sequences in Cluster 0 and 21 sequences in Cluster 1). The amino acid sequence of the corresponding *ftsZ* CDS representing the first four cluster i.e., *Pseudoalteromonas luteoviolacea* (Cluster 0), *Cutibacterium avidum* 44067 (Cluster 1), *Streptococcus agalactiae* 2603V/R (Cluster 2) and *Burkholderia ubonensis* MSMB22 (Cluster 3) were subjected to secondary structure prediction using SSpro module of SCRATCH Protein Predictor (http://scratch.proteomics.ics.uci.edu/)[46]. SSpro catalogues three classes of secondary structure and based on that, the amino acid residues constituting the four FtsZ proteins have been identified as H (alpha helix), E (strand) and C (all the rest secondary structural elements). We have meticulously aligned the *ftsZ* gene sequences with their corresponding amino acid sequence, and secondary structure mark-up sequence generated using SSpro. Using this triple alignment for each of the four representative sequence, we have identified the individual codons coding for each of the different amino acids. Then we have tied the same with the codons encoding the different secondary structures (Figs. 2-5). We have analysed the RSCU values of the *ftsZ* CDS by splitting the sequences according to the tendency of the residues in constituting the three different secondary structural element classes. A graphical representation of the RSCU values is given in Fig. 6. An amino acid wise comparative analysis of the four representative *ftsZ* CDS is discussed in the succeeding sections.

**Table 3:** Clusters of *ftsZ* gene sequences generated using CD-HIT with a similarity threshold of 50 percent.

| Cluster at 50% identity | No. of sequences in the cluster | Representative sequence | Length of the representative sequence |
| --- | --- | --- | --- |
| Cluster 0 | 41 | *Pseudoalteromonas luteoviolacea* | 418 |
| Cluster 1 | 21 | *Cutibacterium avidum* 44067 | 417 |
| Cluster 2 | 9 | *Streptococcus agalactiae* 2603V/R | 419 |
| Cluster 3 | 6 | *Burkholderia ubonensis* MSMB22 | 399 |
| Cluster 4 | 4 | *Butyrivibrio proteoclasticus* B316 | 412 |
| Cluster 5 | 3 | *Acinetobacter johnsonii* XBB1 | 398 |
| Cluster 6 | 2 | *Neisseria gonorrhoeae* FA 1090 | 392 |
| Cluster 7 | 2 | *Geobacter sulfurreducens* PCA | 383 |
| Cluster 8 | 2 | *Anaplasma marginale* str. Florida | 412 |
| Cluster 9 | 1 | *Fusobacterium nucleatum* | 360 |
| Cluster 10 | 1 | *Deinococcus radiodurans* R1 | 371 |
| Cluster 11 | 1 | *Helicobacter pylori* 26695 | 385 |
| Cluster 12 | 1 | *Arcobacter butzleri* RM4018 | 377 |
| Cluster 13 | 1 | *Chromobacterium subtsugae* | 400 |
| Cluster 14 | 1 | *Ralstonia solanacearum* GMI1000 | 408 |
| Cluster 15 | 1 | *Borrelia burgdorferi* B31 | 399 |
| Cluster 16 | 1 | *Selenomonas noxia* ATCC 43541 | 412 |

**Fig 2:**
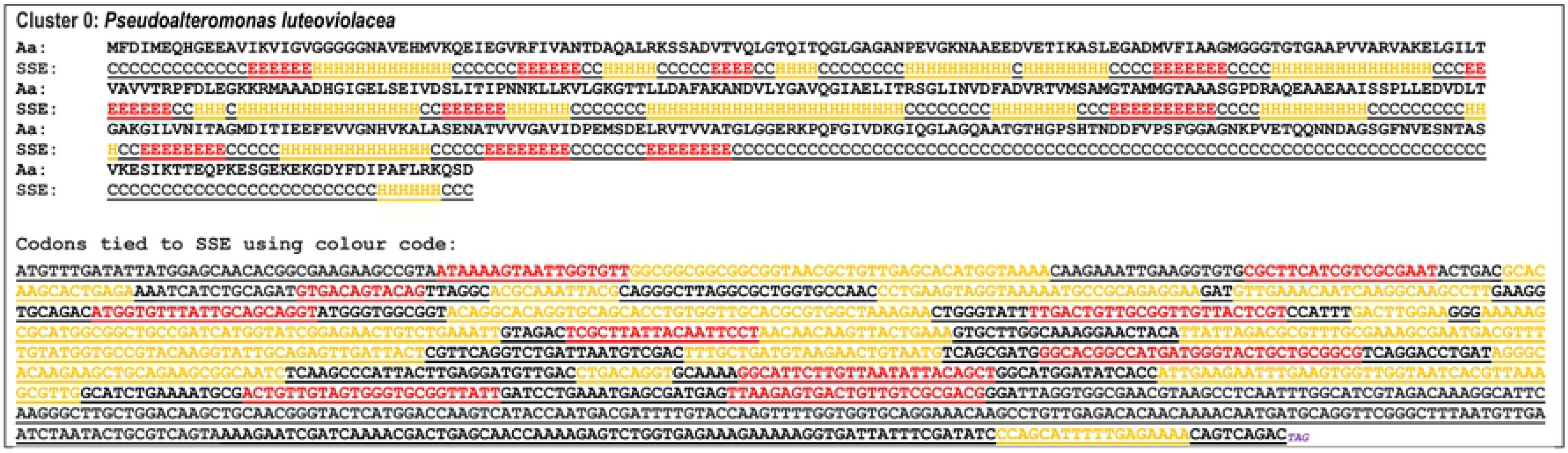
The markup of the FtsZ protein amino acid sequence of *Pseudoalteromonas luteoviolacea* with the secondary structural elements that has been colour coded to tie up with the corresponding codons of the *ftsZ* gene. The codons coding for the *ftsZ* gene has been tied to the secondary structural elements using the following colour code. Orange= residues/codons in helix regions; red= residues/codons in strand regions; black= residues/codons in other secondary structural elements.

**Fig 3:**
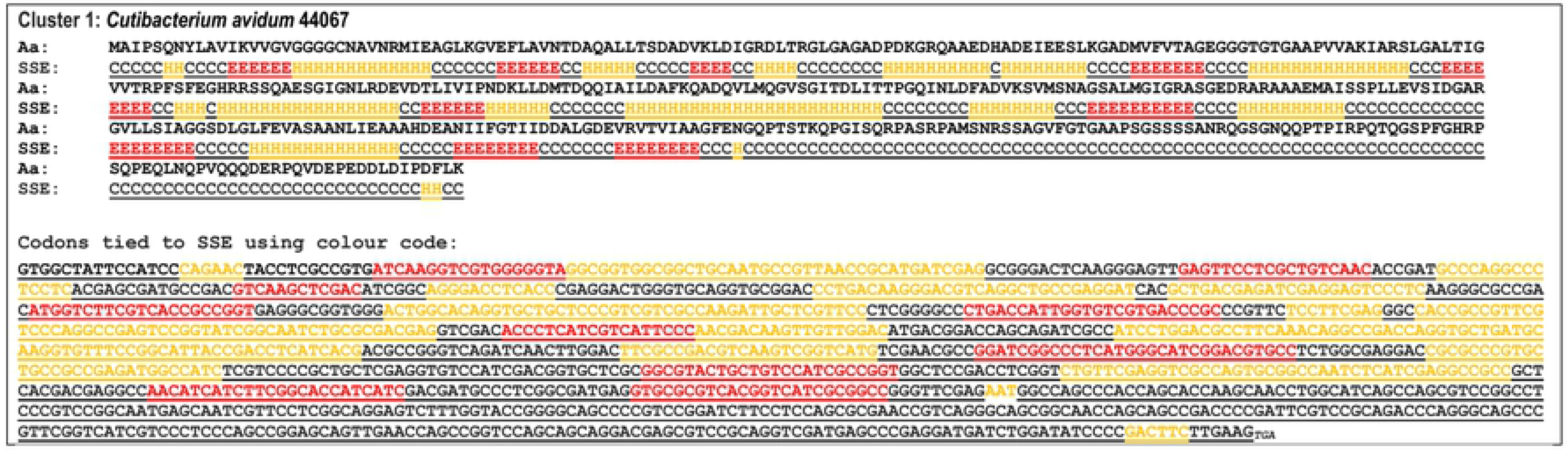
The markup of the FtsZ protein amino acid sequence of *Cutibacterium avidum* 44067 with the secondary structural elements that has been colour coded to tie up with the corresponding codons of the *ftsZ* gene. The codons coding for the *ftsZ* gene has been tied to the secondary structural elements using the following colour code. Orange= residues/codons in helix regions; red= residues/codons in strand regions; black= residues/codons in other secondary structural elements.

**Fig 4:**
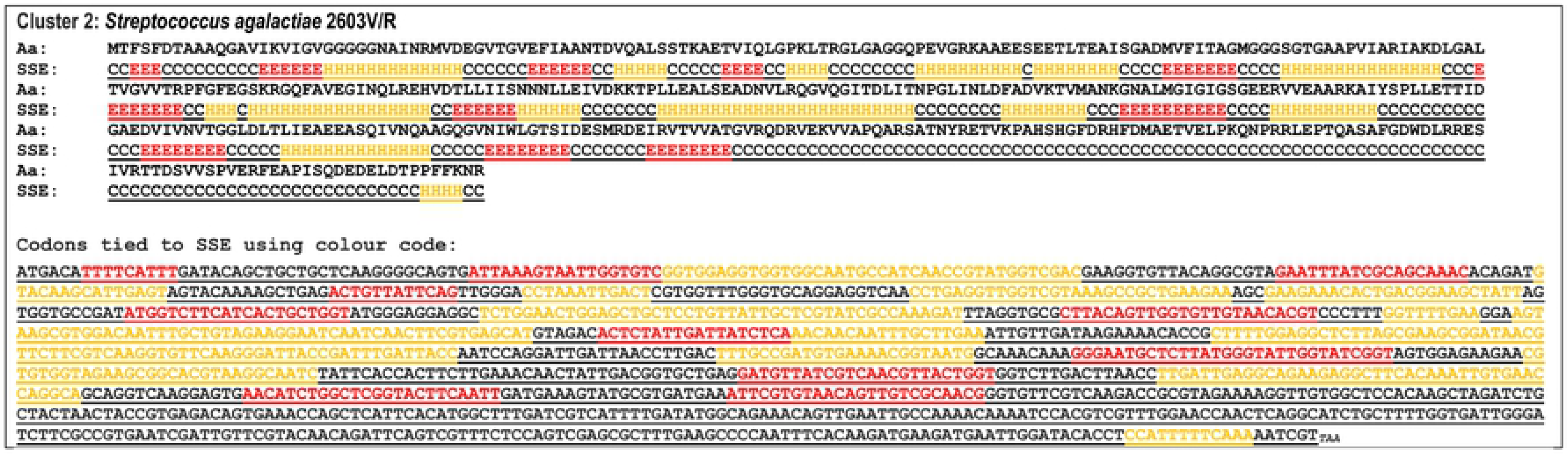
The markup of the FtsZ protein amino acid sequence of *Streptococcus agalactiae* 2603V/R with the secondary structural elements that has been colour coded to tie up with the corresponding codons of the *ftsZ* gene. The codons coding for the *ftsZ* gene has been tied to the secondary structural elements using the following colour code. Orange= residues/codons in helix regions; red= residues/codons in strand regions; black= residues/codons in other secondary structural elements.

**Fig 5:**
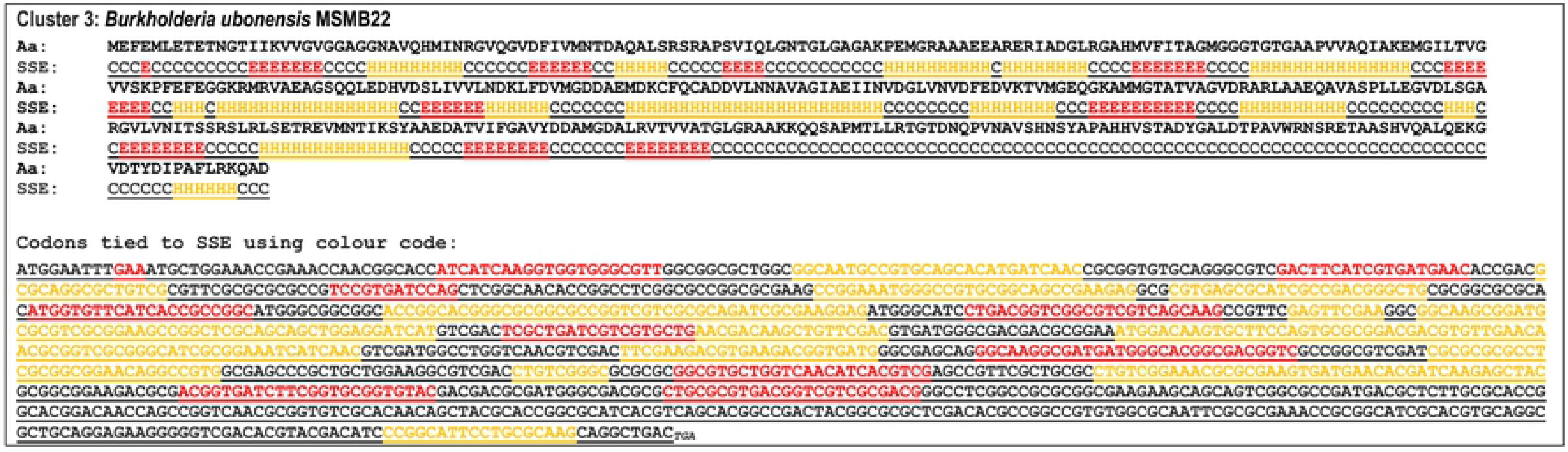
The markup of the FtsZ protein amino acid sequence of *Burkholderia ubonensis* MSMB22 with the secondary structural elements that has been colour coded to tie up with the corresponding codons of the *ftsZ* gene. The codons coding for the *ftsZ* gene has been tied to the secondary structural elements using the following colour code. Orange= residues/codons in helix regions; red= residues/codons in strand regions; black= residues/codons in other secondary structural elements.

**Fig 6:**
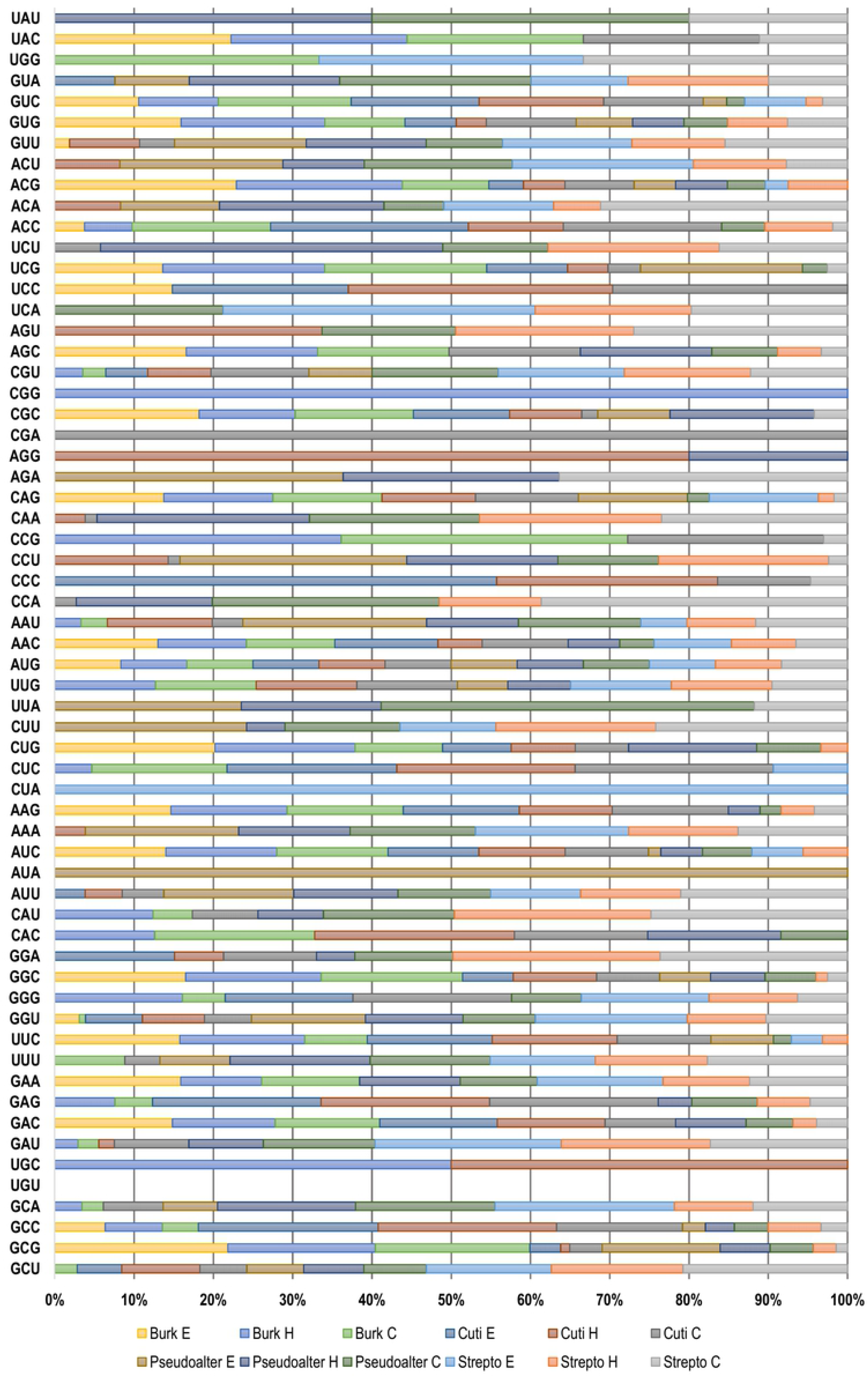
A graphical representation of the relative synonymous codon usage (RSCU) values of the *ftsZ* coding sequences expressed by splitting the sequences according to the tendency of the residues in constituting the three different secondary structural element classes in the four bacterial species. Burk=*Burkholderia ubonensis* MSMB22, Cuti=*Cutibacterium avidum* 44067, Pseudoalter=*Pseudoalteromonas luteoviolacea*, Strepto=*Streptococcus agalactiae* 2603V/R. The suffix H, E and C refers to the helix, strand and other secondary structural elements of the FtsZ protein respectively.

### Amino acid wise comparative RSCU analysis of the helix, strand and other structural element constituting residues

A RSCU analysis of the sense codons used for coding the amino acids of the FtsZ protein was carried out. The triple markup sequences from *Pseudoalteromonas luteoviolacea*, *Cutibacterium avidum* 44067, *Streptococcus agalactiae* 2603V/R and *Burkholderia ubonensis* MSMB22, described in the preceding section was used to classify the codons into three types based on the type of secondary structural elements they constitute. An amino acid wise description of the RSCU of the sixty one sense codons used in structuring of the *ftsZ* CDS is described below. On the basis of chemical nature, the amino acids have been classified into four groups— non-polar, polar basic, polar acidic and polar neutral.

#### Non Polar amino acids

##### Glycine

In case of glycine, the residues constituting the helix in proteins are encoded by the codons GGU, GGA, GGG and GGC. *Burkholderia* does not use the codons GGU and GGA. The codon GGC is used by all the four organisms— *Cutibacterium*, *Burkholderia, Pseudoalteromonas*, and *Streptococcus.* The codon GGU is used by three organisms except *Burkholderia* whereas GGG is shared by *Burkholderia* and *Streptococcus.* GGA is absent only in *Burkholderia.* In case of the strand region, *Cutibacterium* utilizes all the four codons whereas *Burkholderia* and *Pseudoalteromonas* use only two codons GGU and GGC. Likewise, *Streptococcus* also prefers the two codons GGU and GGG only. This suggests that in these three bacterial species there is a preference towards a certain subset of codons in coding the glycine residues positioned in the strand regions. In all the remaining secondary structural elements, all the organisms are found to use GGU, GGG and GGC. The GGA codon was found to be absent in *Burkholderia*.

##### Alanine

The amino acid alanine is near universally encoded by GCU, GCC, GCA and GCG. In the helix region, we observed that *Burkholderia* does not use the codon GCU. GCG and GCC codons are used by all the four organisms. GCA is found to be absent in *Cutibacterium*. All the four codons are found to be employed by *Streptococcus*. But in the strand region, GCU and GCA are used only by *Streptococcus*. Two other codons, GCG and GCC are used by *Burkholderia* alone. In the remaining regions, all the four codons are used randomly by all the organisms.

##### Valine

It is encoded by GUU, GUC, GUA and GUG. In the helix region, GUU is not used by *Burkholderia* but the codon GUG is used by all the four organisms. Codon GUC is found to be absent in *Pseudoalteromonas*, whereas GUA was shared by two organisms, *Pseudoalteromonas* and *Streptococcus*. In contrast to the helix region, in the strand region *Pseudoalteromonas* use all the valine synonym triplets*. Cutibacterium* was found to use three codons, GUG, GUC and GUA, but in *Streptococcus* it was GUU, GUC and GUA. *Burkholderia* majorly uses GUG and GUC and a small frequency of GUU. *Burkholderia* does not use the codon GUA. In all the remaining regions, GUU was not used by *Burkholderia*. All the four organisms use two codons i.e. GUG and GUC. Codon GUA was found only in *Streptococcus* and *Pseudoalteromonas*.

##### Methionine

Since methionine is coded by a single codon AUG, we observed that for all the three regions, the codon AUG is preferred by all the four species.

##### Leucine

It is one of the three amino acids which is encoded by six different codons UUA, UUG, CUU, CUC, CUG and CUA. In the helix elements, the codon CUC is present only in *Burkholderia* and *Cutibacterium*, but CUG is present in all the four organisms. CUU present only in *Pseudoalteromonas* and *Streptococcus.* The codon UUA is found to be used by only one organism– *Pseudoalteromonas.* Codon UUG used by all the organisms whereas the codon CUA is totally absent in the helix region. In the strand region, *Burkholderia* uses only one codon CUG, whereas *Cutibacterium* use the codons CUC and CUG and *Pseudoalteromonas* uses three codons (CUU, UUA, and UUG). *Streptococcus* uses CUA, CUC, CUU and UUG codons. It was observed that there are two codons which are used by only two organisms— CUA is used by *Streptococcus* and UUA by *Pseudoalteromonas* alone. No organism was found to use all the 6 codons. Now if we look at the remaining regions, it was observed that the codon CUA is not used by any of the species. The codon CUG is used by three organisms except *Streptococcus*. Codon CUC is used by *Burkholderia* and *Cutibacterium* whereas CUU and UUA is not used by *Burkholderia* and *Cutibacterium* and *Pseudoalteromonas* does not use the codon UUG.

##### Isoleucine

In the helix regions, the codon AUU is absent in *Burkholderia*. The codon AUC is used by all the organisms whereas AUA remains totally absent. But in the strand regions, codon AUA is used only by *Pseudoalteromonas* which also uses the other two codons AUU and AUC. *Burkholderia* uses only AUC but *Streptococcus* uses AUU and AUC. In the rest of the remaining regions, codon AUU is not used by *Burkholderia.* Similarly AUC is not used by *Streptococcus* but used by the remaining three organisms. Codon AUA is not used by any of the organisms.

##### Proline

In the helix regions, codon CCA is used by only two organisms–*Pseudoalteromonas* and *Streptococcus* whereas codon CCC is used by a single organism, *Cutibacterium.* Three organisms use the codon CCU except *Burkholderia*. CCG is used by *Burkholderia* alone. In the strand regions, out of the four codons of proline, CCC is used by *Cutibacterium* and CCU by *Pseudoalteromonas*. The rest two codons aren’t used. In case of the remaining secondary structural elements, *Cutibacterium* is found to use all the four codons. Codon CCU and CCA are not used by *Burkholderia* whereas the codon CCC is used by *Cutibacterium* and *Streptococcus*; CCG codon is not used by *Pseudoalteromonas*.

##### Phenylalanine

Phenylalanine, a non-polar aromatic amino acid is encoded by two codons– UUU and UUC. Considering the codon usage of the phenylalanine residues in the helix regions, the codon UUU is used by *Pseudoalteromonas* and *Streptococcus* whereas UUC is used by all the organisms except *Pseudoalteromonas.* But in strand elements, codon UUU is only used by *Streptococcus* and *Pseudoalteromonas.* The use of UUC is totally avoided here. In case of the remaining secondary structural elements, UUU codon is used by all the four species.

##### Tryptophan

The amino acid tryptophan is encoded by a single codon UGG in a near universal manner. In case of helix elements of *ftsZ* CDS, this amino acid is totally absent. In the strand elements, UGG is used only by *Streptococcus* whereas in the remaining elements, tryptophan is found to be used by *Burkholderia* and *Streptococcus*.

##### Tyrosine

In the helix regions, we found that the codon UAC is used by *Burkholderia* alone. Similarly *Pseudoalteromonas* use the codon UAU. In strand regions, UAU remains totally absent whereas UAC is used by *Burkholderia* alone. UAC is not used by *Pseudoalteromonas*. In the remaining regions, UAU is found to be used by the organisms *Streptococcus* and *Pseudoalteromonas*.

#### Polar Basic amino acids

##### Histidine

In the helix regions, histidine is coded by CAC in three of the organisms except *Streptococcus*. Similarly, another codon CAU is preferred in the helix regions buy all the three organisms except *Cutibacterium.* In the extended strand regions, our analysis shows that the amino acid histidine isn’t used by any of the four organisms. For the rest of the remaining secondary structural elements CAU is preferred by all the four organisms except *Streptococcus* which uses CAC.

##### Lysine

This amino acid is encoded by two codons— AAA and AAG. In the helix regions lysine is found to be coded by the homo triplet AAA in the studied organisms except *Burkholderia*. AAG was found to be employed by all the four organisms. In the strand regions, the triplet AAA is used by the organisms *Pseudoalteromonas* and *Streptococcus* whereas AAG is preferred by *Burkholderia* and *Cutibacterium.* For the remaining secondary structural elements, the preference for AAA is restricted to *Pseudoalteromonas* and *Streptococcus*, a scenario exactly similar to the strand region.

##### Arginine

This is one of the three amino acid which is encoded by the maximum number of degenerate codons – CGU, CGC, CGA, CGG, AGA and AGG. In the helix regions, none of the four organisms were found to use the codon CGA. The remaining organisms display preference towards the use of specific codons. The codon AGA is used by only one organism *Pseudoalteromonas* whereas AGG is preferred by *Cutibacterium* and *Pseudoalteromonas*. *Streptococcus* does not use the codon CGC whereas CGG is used by *Burkholderia* alone. *Streptococcus* uses the codon CGU in the maximum frequency than the remaining organisms whereas it was found to be absent in *Pseudoalteromonas.* In the strand regions only three codons are used out of the six– AGA, CGC and CGU. This suggests the preference of the organism towards specific codons for encoding the amino acids that have the propensity to be included in the strand regions of FtsZ protein. AGA is used by *Pseudoalteromonas* alone whereas CGC is used by all except *Strepptococcus. Burkholderia* does not use the codon CGU. *Cutibacterium* on the other hand uses two codons– CGC and CGU whereas *Streptococcus* use only CGU. *Burkholderia* uses the codon CGC only for encoding the amino acids in the strand regions.

In the remaining structural elements, out of the six codons two are totally absent and this are CGG and AGA. CGA is used only by *Cutibacterium*, whereas AGG is used only by *Streptococcus.* The codon CGC is used by three organisms except *Pseudoalteromonas.* CGU is found to be used by *Cutibacterium*, *Pseudoalteromonas, Streptococcus* and comparatively in lesser frequency by *Burkholderia*.

#### Polar acidic amino acids

##### Aspartic acid

In the helix region, all the four organisms use both the codons GAU and GAC, but the frequency of GAU used by *Cutibacterium* is very low. In contrast to the helix regions, in strand regions we found that *Pseudoalteromonas* does not use aspartic acid. *Streptococcus* use GAU alone whereas GAC is used by *Burkholderia* and *Cutibacterium*.

##### Glutamic acid

This amino acid is represented by the codons GAA and GAG. In the helix, GAG is used by all the four organisms whereas GAA is used by all except *Cutibacterium*. In the E region, GAA is used by *Burkholderia* and *Streptococcus* whereas GAG is used by *Cutibacterium* alone. Both the codons are found to remain absent in *Pseudoalteromonas.* In the rest of the structural elements, GAG is preferred by all the organisms, but GAA is not used by *Cutibacterium*.

#### Polar Neutral amino acids

##### Serine

It is encoded by six codons— UCU, UCC, UCA, UCG, AGU, AGC. We have observed a preferential usage of certain codons encoding the different amino residues constituting the different structural elements. In the helix regions, AGC is used by all the organisms except *Cutibacterium*. The codon AGU is found to be preferred by *Streptococcus* and *Cutibacterium*. UCA and UCC codons are found to be used only by *Streptococcus* and *Cutibacterium* respectively. *Burkholderia* and *Cutibacterium* was found to prefer UCG, whereas *Pseudoalteromonas* and *Streptococcus* use UCU. The use of the codon UCU by *Pseudoalteromonas* was found to be comparatively higher than the rest of the organisms. In the strand regions, the codon AGU and UCU were found to be avoided by all the four organisms. *Burkholderia* prefers the codons AGC, UCC and UCG whereas *Cutibacterium* prefers only two codon UCC and UCG. *Pseudoalteromonas* and *Streptococcus* was found to use only one codon which is UCG and UCA respectively. In the rest of the structural elements, AGC was found to be preferred by all the four organisms. The codons AGU and UCA are used by *Streptococcus* and *Pseudoalteromonas* whereas UCC is used by *Cutibacterium* alone. All the four organisms use the codon UCG, but in *Burkholderia* the frequency of usage is relatively greater.

##### Threonine

In the helix elements, ACC and ACA are preferred by three organisms. ACC remains absent in *Pseudoalteromonas* whereas ACA is absent in *Burkholderia.* All the four organisms preferentially use the codon ACG but ACU is absent only in *Burkholderia.* In the extended strand elements, ACG used by all the organisms. ACA and ACU codons are used by *Streptococcus* and *Pseudoalteromonas* whereas *Burkholderia* and *Cutibacterium* use the codon ACC.

##### Asparagine

This amino acid is encoded in general by two codons– AAU and AAC. In the helix regions, AAC is preferred by all the organisms. Likewise codon AAU is also used by all the organisms but the relative usage frequency is very low in *Burkholderia.* But in the strand region, AAU is used by two organisms *Pseudoalteromonas* and *Streptococcus*. The codon AAC is employed by all the organisms except *Pseudoalteromonas.* In the remaining structural elements, we did not observe any fixed preference for a particular codon in the organisms considered in this analysis.

##### Glutamine

In case of helix regions, codons CAA and CAG are used by three organisms. CAA was absent in *Burkholderia* whereas CAG was absent in *Pseudoalteromonas*. The use of the amino acid glutamine in the helix region was absent in *Cutibacterium*. CAG is used by three organisms (*Burkholderia, Pseudoalteromonas and Streptococcus*) but not used by *Cutibacterium*. The codon CAA was not used by any of the organisms in the strand regions. In the other secondary structural elements, the codon CAG is used by all the organisms whereas the codon CAA is used by all the organisms except *Burkholderia*.

##### Cysteine

In the helix elements, the codon UGU was avoided by all the organisms. The use of this sulphur containing amino acid in the helix regions of *Burkholderia* and *Cutibacterium* are found to be fulfilled by the codon UGC. Apart from the helix structural elements, cysteine was found to be totally absent in the other secondary structural elements in all the four organisms.

Our study clearly shows that a differential RSCU pattern is evident in the coding nature of the various secondary structural elements of the FtsZ proteins from different bacteria. It may be suggested that this variation could be attributed to the differential folding pattern of the different domain region of the FtsZ protein. The FtsZ protein has two major domains— one is the GTPase domain and the other is the C-terminal domain. Our findings suggest that the use of specific codons coding for the amino acids in the different secondary structural elements of the FtsZ protein is less organism specific but more codon specific. The helix regions demonstrates a comparative higher bias towards use of specific codons in coding the amino acids than the strand or the other secondary structural element regions.

## Conclusions

The FtsZ protein is ubiquitous in bacteria and plays a vital role in bacterial cell division. From the evolutionary stand point it might be regarded as the counterpart of the eukaryotic tubulin protein. Our study of the gene sequences coding for FtsZ from 142 bacterial species demonstrating different lifestyle and Gram nature showed that the degree of sequence identity among the protein fluctuates from a mere thirteen percent to a whooping ninety eight percent. This is suggestive of a compositional variability both in the coding sequence and amino acid sequence. We found that about one third of the selected organisms depicted more than ten percent GC variation in their *ftsZ* CDS compared to their genomic GC content. Thus, our study clearly suggest that the nucleotide composition of the gene coding for FtsZ protein in a large number of species deviates significantly from their genomic GC content. The codon usage pattern analysis also demonstrated that the *ftsZ* gene of about ninety three percent of the organisms showed relatively biased codon usage profile. In this study, we have also captured the existence of a ‘core’ set of codons in the structuring of the *ftsZ* gene despite the presence of a varying degree of identity among the *ftsZ* sequences. This is probably due to the constraint exerted by nature to maintain form and function in an important physiological protein FtsZ that plays a major role role in successful completion of bacterial cell division. By the utilization of inferential statistical methods such as a two way ANOVA, we were able to capture the influence of Gram nature of the bacteria and their lifestyle pattern on the amino acid compositional frequency of the FtsZ protein. Finally, a cluster analysis followed by an amino acid wise comparative RSCU analysis of the different secondary structural elements of the FtsZ protein tied with the *ftsZ* CDS, demonstrated the presence of bias towards specific triplet codons coding the amino acids of the different secondary structural elements of a multi domain protein like FtsZ. In conclusion, it may be stated that the *ftsZ* gene coding for an indispensable cell division protein called FtsZ in a large number of bacteria, differing in terms of cellular morphology, physiology, biochemistry and a host of other features displays a very biased codon usage pattern with a highly skewed GC content. Along with the existence of a preferred ‘core’ set of codons, the different secondary structural elements of the multi-domain FtsZ protein was also found to display bias towards specific synonymous codons particularly in the helix and strand regions. All these suggest that in an indispensable and vital protein such as FtsZ, there is an inherent tendency to maintain form and structure for optimized performance in spite of the extrinsic variability in coding features.

## Acknowledgements

The authors are grateful to Prof. Subhasis Mukhopadhyay and Late Prof. A. K. Bothra for their support and encouragement.

